# Cell-specific heteroclinic orbits govern interaction dynamics in inertial microfluidics for circulating tumour cell separation

**DOI:** 10.64898/2026.09.15.751739

**Authors:** Roslyn Hay, Chiara Ghera, Jian Zhou, Ian Papautsky, Timm Krüger, Benjamin Owen

## Abstract

Cancer survival rates increase with earlier diagnosis. However, many cancers are diagnosed only after symptoms develop, in the later stages of cancer progression. We need a diagnostic tool that can detect cancer earlier. Circulating tumour cells (CTCs) are cells that detach from the main tumour and can enter the bloodstream. The capture and analysis of CTCs can provide an early indicator of cancer. However, rarity of CTCs in blood makes their efficient capture challenging. Inertial microfluidics can be utilised for a label-free separation of CTCs from white blood cells (WBCs) by manipulating cells in the microchannel based on cell properties, achieving good separation performance. Inertial microfluidic devices mainly rely on size-based separation. However, residual WBC carryover limits complete separation. Here, we show that cell mechanical heterogeneity plays a large role in the separation process and should be considered in the design of these devices. We simulate the migration dynamics of CTCs and WBCs in a straight microchannel using a 3D lattice-Boltzmann-immersed-boundary-finite-element solver. Our results demonstrate that WBC migration behaviour changes depending on the deformability of a CTC. The size and deformability of the cells have been shown to determine the cell-specific heteroclinic orbits leading to different interaction types, these interactions alter WBC migration resulting in more/less WBC carryover. This work highlights that both cell-specific heteroclinic orbits and single cell migration rates should be considered in the design of inertial microfluidics for separation.

## 1 Introduction

Cancer accounts for 1 in 6 deaths each year, resulting in nearly 10 million deaths worldwide in 2024^2^. Current detection methods generally require prior knowledge of cancer location, which is usually obtained only after symptoms develop. Circulating tumour cell (CTC) capture and analysis may provide clinically useful information through minimally invasive blood samples. CTCs are cells that detach from a primary or metastatic tumour and enter the blood. However, CTC isolation remains challenging since CTCs are rare, typically occurring at concentrations of approximately 5-100 cells per ml of blood, and exhibit substantial heterogeneity in cell size and deformability due to cancer type and stage ^3,4^.

Inertial microfluidics provides a promising approach for the high-throughput, label-free separation of CTCs from blood samples ^1,5–7^, based on cell properties, device geometry, and flow conditions ^8^. These devices use fluid inertia to direct suspended cells along size-dependent migration pathways toward predictable cross-sectional positions ^8 9^. Numerical modelling has played an important role in elucidating the mechanisms governing inertial migration and in supporting the design of increasingly complex inertial microfluidic systems ^10^.

An established approach exploits differences in cell size to separate CTCs from white blood cells (WBCs). In the device developed by Zhou et al. ^6^, larger CTCs migrate laterally towards the target stream more rapidly than smaller WBCs within the straight channel, allowing the channel length to be selected such that CTCs enter the target outlet while WBCs ideally remain within the waste streams.

This platform was recently applied to CTC isolation from blood samples from patients with PDAC. ^11^ In that study, the measured size distributions of PANC1 cells and WBCs supported the physical basis for size-dependent separation, and representative outlet measurements demonstrated preferential routing of the larger PANC1 cells to the target while WBCs remained enriched in the waste outlets. Nevertheless, residual WBCs in the target stream remain an important concern because contaminating cells can interfere with downstream analysis of enriched CTC samples.

The conventional description of this separation process therefore centres on differences in single-cell migration rates: larger cells generally migrate more rapidly towards their equilibrium positions than smaller cells ^12 13^. However, separation occurs within a suspension, rather than between isolated cells, and cell–cell interactions can perturb these migration trajectories. Previous work has shown that size heterogeneity influences particle–particle interactions, while differences in particle deformability can also affect the formation of particle pairs. These observations suggest that single-cell migration behaviour alone is insufficient to describe the separation of mechanically heterogeneous CTC and WBC populations. Instead, understanding WBC migration toward the target stream requires consideration of both the baseline migration behaviour of individual cells and the interactions that occur between them ^14^.

Inertial migration can be described in terms of the heteroclinic orbit (HCO) followed by each suspended cell or particle. Cell migration towards equilibrium occurs in two stages: a comparatively rapid migration towards the HCO, followed by slower migration along the HCO towards the stable equilibrium position ^15^. Therefore, HCOs describe the cross-sectional pathways occupied by cells over a substantial part of their downstream migration. Their locations and shapes depend on particle properties, including size and deformability, as well as channel geometry and flow conditions. Previous work has shown that particle size influences both migration rates and migration pathways, while more deformable particles migrate closer to the channel centre than less deformable particles ^16^. This raises an important consequence for heterogeneous suspensions: cells with different mechanical properties may follow distinct cell-specific HCOs, and the spatial separation between these pathways may determine how frequently, and for how long, different cell types interact. Moreover, interactions may displace cells from the trajectories predicted from their isolated migration behaviour. This is particularly relevant for CTC separation because patient-derived CTCs vary not only in size but also in their mechanical properties. WBCs similarly exhibit variation in size and deformability across different subpopulations. ^17,18^

Although deformability has been shown to affect inertial migration and particle pairing, it remains unclear how mechanical heterogeneity alters the spatial relationship between CTC and WBC migration pathways and, consequently, their interaction dynamics. In particular, it is not known whether proximity between the cell specific HCOs promotes longer CTC-WBC interactions or whether such interactions produce net WBC migration toward the target stream. We therefore hypothesise that CTC and WBC deformability modifies the separation between cell-specific HCOs of CTCs and WBCs, altering the nature and duration of CTC–WBC interactions and thus changing WBC migration toward or into the target stream.

In this work, we investigate how cell deformability alters the relative migration pathways of CTCs and WBCs and the interactions that occur between them in an established straight channel inertial microfluidic device. Using 3D numerical simulations, we first determine the HCOs and migration rates of isolated CTCs and WBCs and then evaluate interaction-mediated departures from these baseline trajectories in mixed cell suspensions. We show that mechanical heterogeneity alters the CTC and WBC HCOs and is associated with distinct short and long interaction. Reduced separation between the cell-specific HCOs increases the occurrence of long CTC-WBC interactions, which produce greater net WBC migration toward the target stream than short encounters. These findings demonstrate that inertial microfluidic separation cannot be understood from single-cell migration rates alone. In addition to controlling engagement with cell-specific migration pathways, device analysis must account for the proximity pathways followed by target (CTC) and background (WBC) cells and the interactions that this proximity enables.

## 2 Methods

### 2.1 Physical model and microfluidic set-up

Fig. 1a shows the previously reported inertial microfluidic device, in which two lateral inlets carrying lysed blood sample and one central inlet carrying buffer solution merge into a straight channel. After traversing the straight channel, the cells exit through one central target outlet and two lateral waste outlets. Details of device fabrication and experimental operation were reported previously ^1,6,11^.

**Fig. 1.**
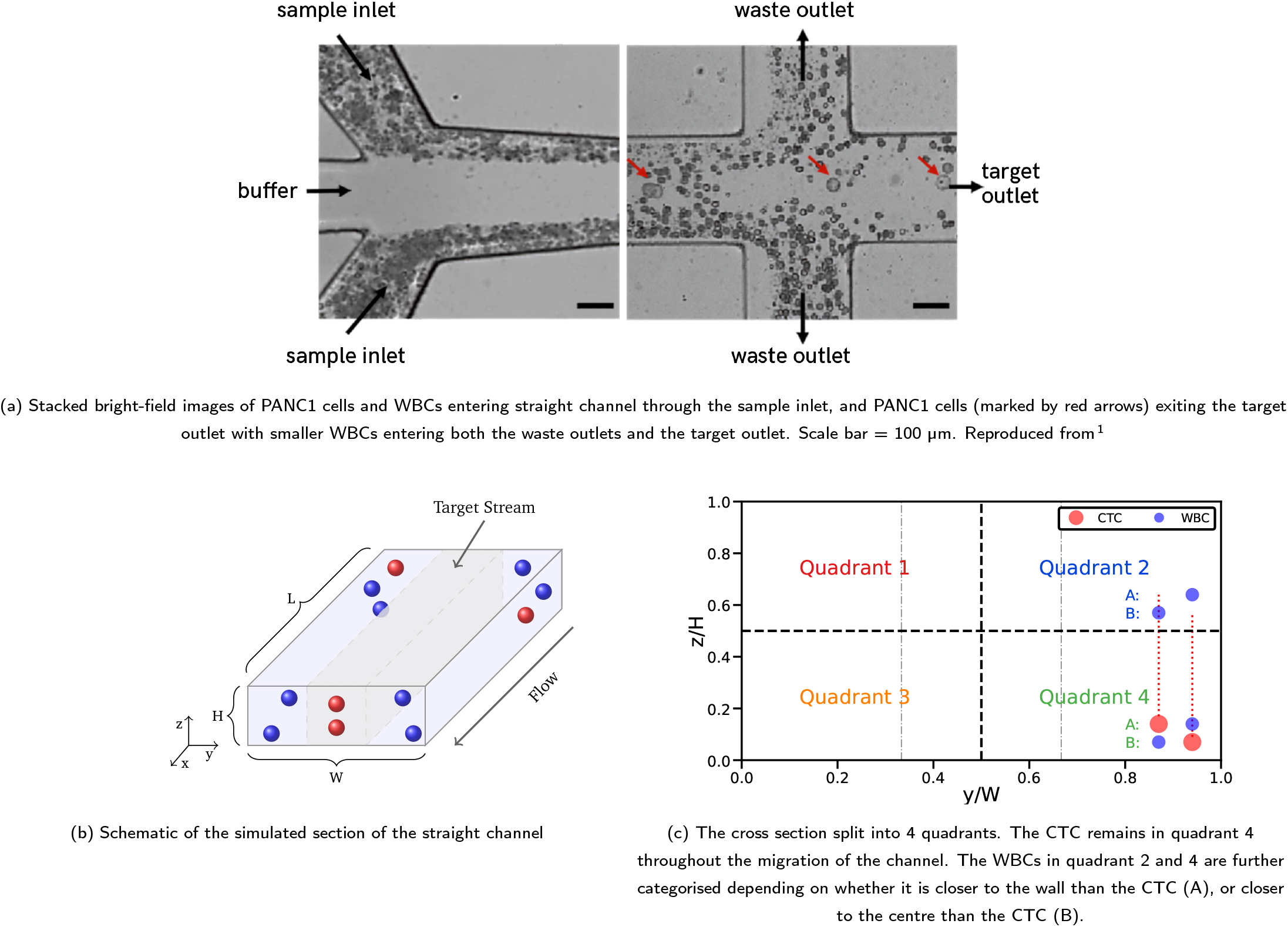
Inertial microfluidic device design. (a) Inlets containing the lysed blood sample, connected to the buffer inlet, form the straight channel. AIM: CTCs migrate to and leave via the target stream and WBCs leave via the waste streams. (b) Simulation domain of the Inertial Microfluidic Device. Cells are initialised in one of the four quadrants in the cross section, outside the target stream, shown in (c). (d) Experimental image of the WBC carryover problem.

We considered a straight channel, with a rectangular cross-section, to represent the straight section of the device, as seen in Fig. 1b. The dimensions of the straight channel in Fig. 1a were length *l* = 30 mm, width *W* = 150 *µ*m, and height *H* = 50 *µ*m.

The coordinate system was defined such that 0 ≤ *y* ≤ *W* and 0 ≤ *z* ≤ *H*, and the flow was along the *x*-axis. A small segment along the length of the channel was modelled and flow-wise periodic boundary conditions were used such that our simulation domain had length *L* = *l/*80. The fluid and cells that exited the domain at *x* = *L* re-entered the domain at *x* = 0, and the accumulative downstream positions of the cells were recorded over time. We ensured the simulated channel length was sufficiently long so that periodic images of cells did not interact with each other, validated in ^14^.

The buffer and inlet fluids were modelled as being incompressible, having the same Newtonian kinematic viscosity *ν* and density *ρ*. The fluids were treated as miscible, such that there are no surface tension effects. Following previous work ^14^, the two sample inlet flow rates and the central buffer flow rate were assumed to be equal, corresponding to the 1:1:1 sample:buffer:sample flow split. The total flow rate was denoted as *Q*. The study examined only the 1:1:1 baseline operating condition, while the effects of varying the inlet flow split were investigated separately in a related study.

The cells were modelled as deformable, neutrally buoyant capsules that were spherical in their undeformed state and contained the same fluid as the surrounding buffer. The capsule membrane was modelled as a thin hyperelastic material governed by the Skalak model ^19^:

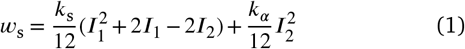

where *w*_s_is the areal energy density, *k*_s_and *k*_*α*_are the elastic shear and area dilation moduli, and *I*_1_ and *I*_2_ are the in-plane strain invariants ^20^. We include a membrane bending energy

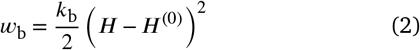

where *k*_b_is the bending modulus, and *H* and *H*^(0)^ are the trace of the surface curvature tensor and the spontaneous curvature, respectively.

The CTCs and WBCs were coupled to the fluid dynamics, with the no-slip condition holding at the surface of the cells, and the cells interacted hydrodynamically with each other.

The diameters of the two cell types were *d*_CTC_= 17 *µ*m and *d*_WBC_= 11 *µ*m, corresponding to the CTC and WBC diameters, respectively. These values were selected based on prior measurements. Unless otherwise specified, in our system, the suspension consisted of eight WBCs and one CTC, equivalent to a cell concentration of 0.2%. The cells were initialised randomly within the two waste streams in one of the four quadrants Fig. 1c, corresponding to 0 ≤ *y*_0_≤ *W /*3 and 2*W /*3 ≤ *y*_0_≤ *W*.

### 2.2 Characteristic scales, dimensionless groups and relevant parameters

We defined the flow characteristics using the channel Reynolds number:

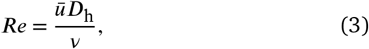

where *ū* is the average velocity in the channel and *D*_h_= 2*W H/*(*W* + *H*) is the hydraulic diameter of the channel. Throughout this work, we kept *Re* = 50 fixed.

We used a dimensionless number to quantify the deformability of the cells, defined as the Laplace number

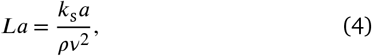

where *a* is the respective cell radius and *k*_s_is the shear modulus. Compared to the capillary number, the Laplace number only depended on material properties and, therefore, was independent of flow conditions. Note that increasing the Laplace number corresponds to decreasing the deformability of the cell. Different degrees of cell deformability were considered by using *La* = [10, 20, 30, 50, 100]. We varied the Laplace number parameter in our simulations by changing the shear modulus, found in Tab. A2. The area dilation modulus and the bending modulus also varied due to the deformability of the cell, as we non-dimensionalised by the shear modulus such that 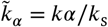 and 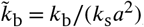, respectively.

Other relevant non-dimensional groups were the confinement of each cell type,

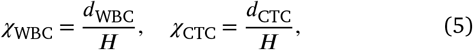

and channel aspect ratio,

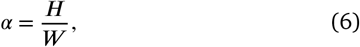

which in the current work are *X*_WBC_= 0.22, *X*_CTC_= 0.34 and *α* = 1*/*3.

A list of the physical parameters alongside their numerical counterparts, as well as the dimensionless parameters are given in Tab. A1. The parameters varied for different cell deformability are provided in Tab. A2. In our results, the particles’ positions were normalised either by the dimensions of the cross sections or the CTC diameter, unless otherwise stated.

### 2.3 Numerical Methods

An in-house BioFM code was used to simulate the fluid flowing through the channel and model the CTCs and WBCs ^20^. This utilised the lattice Boltzmann (LB) method to simulate the fluid, the finite element method (FEM) for particle dynamics and the immersed boundary method (IBM) for fluid-structure interaction. Validation of this code with deformable particles in inertial flow has been demonstrated previously ^16,21^. Here, we provide an overview of the model.

The LB method utilised a D3Q19 lattice together with the BGK collision operator with relaxation time *τ* and Guo’s forcing scheme that applied a constant body force to drive flow. The kinematic viscosity of the fluid was related to *τ* according to

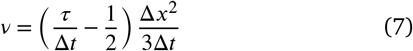

where Δ*t* is the time step and Δ*x* is the lattice spacing. To enforce the no-slip condition on the inner surface of the channel, a half-way bounce-back method was implemented. Periodic boundary conditions were applied in the flow-wise direction.

The FEM was employed to discretise the surface membrane of the CTCs and WBCs by *N*_*f*_flat triangular faces represented by three nodes, such that neighbouring nodes have an average distance of the grid space, Δ*x*. The dynamics of the cells and the fluid were coupled using the IBM with a three-point stencil. The fluid’s velocity was interpolated at the cell nodes, while forces, arising from the cell’s deformation, were spread back to the fluid; this satisfies the no-slip condition between the cell’s surface and the fluid. Particle repulsion models were unnecessary because the suspension is dilute, hydrodynamic forces were sufficient to ensure that particles do not come into contact with each other or the wall with a fluid layer of at least 2Δ*x* separating them.

### 2.4 Classification of short and long CTC-WBC interactions

The long interactions between a CTC is and WBCs was hypothesised to alter WBC migration and thus affect the separation. Because stable CTC-WBC pairs were not expected to persist over the full channel length, interactions were classified according to cell proximity, relative axial velocity, and duration. We defined a long interaction using two criteria:

1. The CTC and WBC remained within an axial distance of 4*d*_CTC_when the WBC was located in quadrant 2 or 4, or 6*d*_CTC_when the WBC was located in quadrant 1 or 3, for 3000 time steps. The 3000 time step threshold corresponded to ≈ 117*µm* downstream distance.
2. During the same interval, the normalised axial velocity difference, 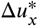, between the two cells was less than the specified tolerance *ϵ* = 0.03.

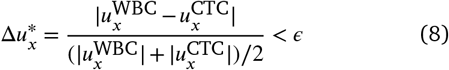

An interaction was classified as short when the CTC and WBC satisfied the corresponding axial proximity criterion but did not satisfy both the duration and relative axial velocity criteria required for a long interaction.

### 2.5 Mean separation between cell-specific HCOs

The separation distance between the HCOs of a CTC and a WBC was quantified using an area-based mean separation metric:

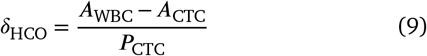

where *A*_WBC_and *A*_CTC_are the cross-sectional areas enclosed by the WBC and CTC HCOs, respectively, and *P*_CTC_is the perimeter of the CTC HCO. This metric represents the area enclosed between the two HCOs normalized by the perimeter of the CTC HCO. Smaller values of *δ*_HCO_indicate more closely aligned cellspecific migration pathways.

## 3 Results and Discussion

### 3.1 CTC and WBC migration in mixed suspensions

We first investigated the effect of CTC deformability (*La*_CTC_= 10 and 100) on the migration of CTCs and WBCs. The deformability of the WBCs was kept constant in a less deformable state (*La*_WBC_= 100); as generally WBCs are less deformable than CTCs. ^17^ A WBC-only case was included as a reference. In each simulation, eight WBCs were placed randomly within the domain. Eight distinct WBC initial-position configurations were generated and used consistently across all cases, including the WBC-only case and the cases containing a CTC. When present, the CTC was initialised at the same position in every configuration. This matched design allowed differences in WBC migration among cases to be attributed to the presence and deformability of the CTC rather than to differences in the initial WBC distribution. The total volume fraction in the CTC-containing suspensions was 0.2%, representing a dilute suspension used with the real-world device.

Fig. 2 compares the combined cell trajectories from the eight matched initial-position configurations. CTC trajectories are shown in red and WBC trajectories in grey. Trajectories are displayed in a top-down *x*−*y* plane view, corresponding to the viewing plane used in experiments. The thick dashed lines indicate the boundaries of the geometrically defined target stream, and the thin dashed line highlights the channel centreline.

**Fig. 2.**
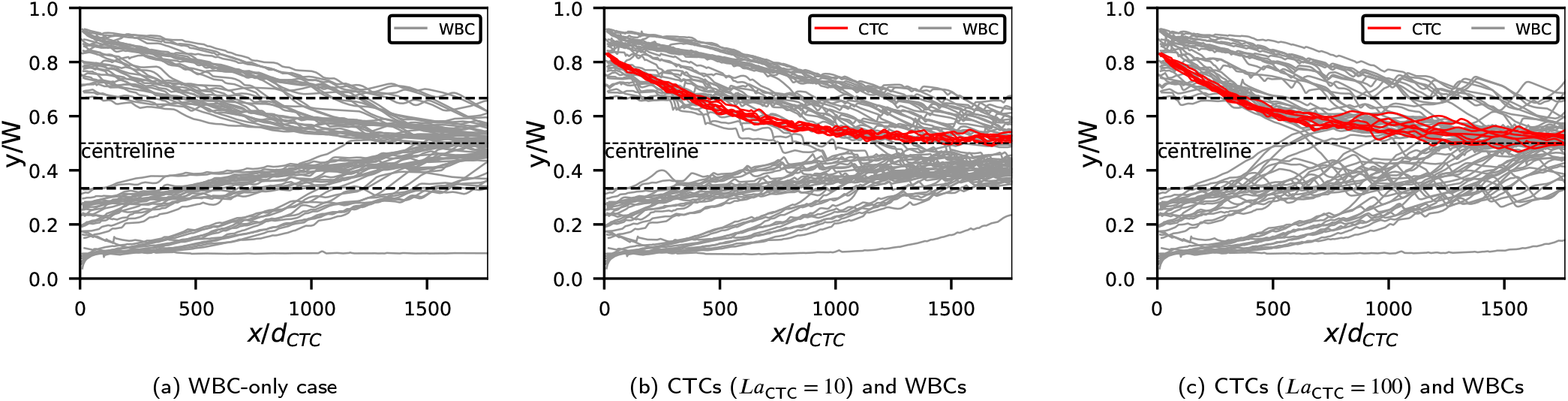
Top-down view of stacked trajectories from eight separate configurations of three cases: (a) WBC-only case, totalling 64 cell trajectories; (b) CTCs and WBCs for *La*_CTC_ = 10, totalling 72 cell trajectories; (c) CTCs and WBCs for *La*_CTC_ = 100, totalling 72 cell trajectories. In all cases, WBCs have the same deformability (*La*_WBC_ = 100). The thick and thin dashed horizontal lines indicate the target stream partition and the channel centreline, respectively.

In the WBC-only case, WBCs first reached the centreline at *x/d*_CTC_≈ 1100 (Fig. 2a). Thus, the upstream portion of the target stream remained free of WBC trajectories. Delayed WBC entry into this stream is desirable because WBCs constitute contaminants in the collected CTC fraction. When a CTC was present (Fig. 2b and 2c), WBCs entered the target stream earlier and first reached the centreline at *x/d*_CTC_≈ 800 for *La*_CTC_= 10 and at *x/d*_CTC_≈ 500 for *La*_CTC_= 100. Thus, relative to the WBC-only case, the first observed centerline arrival occurred ≈ 300*/d*_CTC_earlier in the presence of the more deformable CTC and 600*/d*_CTC_earlier in the presence of the less deformable CTC.

CTCs that are less deformable (*La*_CTC_= 100, Fig. 2c) were as-sociated with a greater variation in WBC migration rates than the more deformable CTCs (*La*_CTC_= 10, Fig. 2b). Because trajectory slope provides only a qualitative indication of migration rate, the CTC-induced changes in WBC migration were quantified in the subsequent section. These results show that the presence and deformability of a CTC altered WBC migration relative to the WBC-only reference. However, these trajectories do not reveal whether the differences arose from changes in isolated-cell migration or from CTC-WBC interactions. We therefore next examined the baseline behaviour for migration of each cell type in isolation.

### 3.2 Behaviour of single CTC and WBC

Single-cell behaviour was studied to establish the migration expected in the absence of cell-cell interactions. We initially focused on the second stage of migration, during which cells migrate along the HCO, because this stage was slower and accounted for most of the downstream migration within the channel. Furthermore, migration along the HCO was independent of initial position once a cell had reached its HCO ^14^.

The HCOs for CTCs and WBCs with *La* = [10, 100] are shown in Fig. 3a and additional HCOs for *La* = [20, 30, 50] are provided in Fig. A1. The HCOs were obtained by initialising single cells near the channel wall at the farthest point from their respective equilibrium positions and allowing the cells to migrate all the way to their equilibrium positions. All HCOs had a similar overall shape. The CTCs, which were larger than the WBCs, followed HCOs located closer to the channel centre than those followed by the WBCs, consistent with previous findings ^14^. Cells that are more deformable also followed HCOs located closer to the channel centre ^22^. As a result, a more deformable WBC (*La*_WBC_= 10) and a less deformable CTC (*La*_CTC_= 100) followed closely aligned HCOs.

**Fig. 3.**
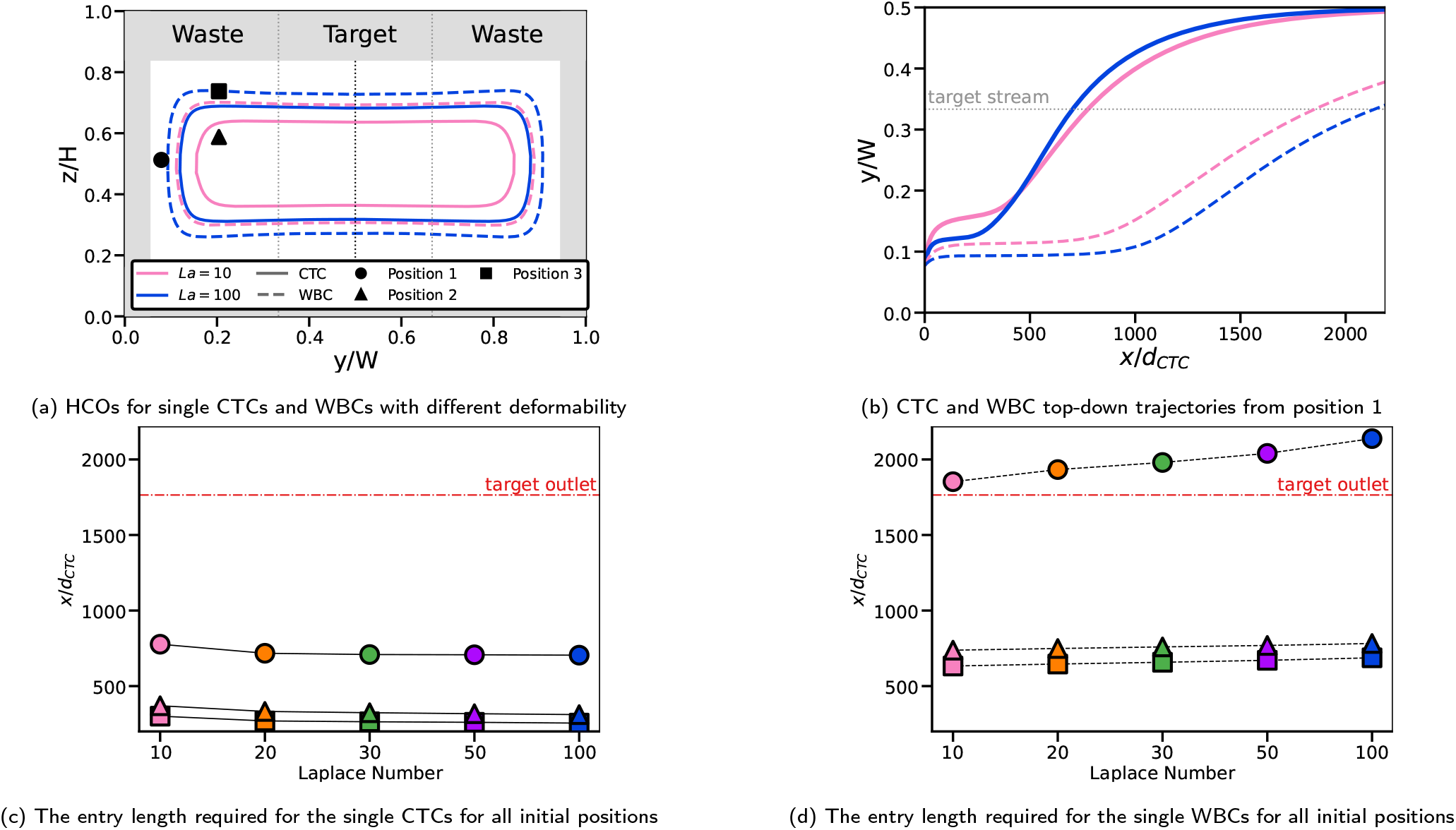
Behaviour of single CTC (solid lines) and WBC (dashed lines) with varied deformability. (a) HCOs for the CTC and WBC with *La* = 10, 100. Markers represent three different initial positions; position 1 (circle), position 2 (triangle), and position 3 (square). (b) Top-down trajectories for a single CTC and WBC, with *La* = 10, 100, for initial position 1, denoted in (a). Target stream partition marked by dotted line. Axial entry length required for CTCs (c) and WBCs (d) to cross into target stream for all initial positions and *La* = 10, 20, 30, 50, 100. Target outlet length marked by dash-dot line.

We next examined the effect of initial position using three distinct initial positions, shown in Fig. 3a, that represented possible locations within and outside the investigated HCOs. Position 1 represented an extreme case near the horizontal midplane with the largest investigated distance between initial and equilibrium positions. Positions 2 and 3 had the same horizontal coordinate but were located within and outside all investigated HCOs, respectively.

Fig. 3b shows the trajectories of cells initialised at position 1 for values *La* = [10, 100] and Figs. [3c–3d] show the axial distance required for the CTCs and WBCs to enter the target stream for all three initial positions and all investigated values of *La*. Cell size had confinement-dependent differences in the entry length to the target streams, with the larger CTC (solid lines) entering the target stream earlier than the smaller WBCs (dashed lines) in all investigated scenarios. This result was consistent with the size-dependent operating principle for the microfluidic device. Initial position also strongly affected the entry distance, consistent with previous work ^16^. Cell initialised at position 1 required a substantially greater downstream distance to enter the target stream than cells initialised at positions 2 and 3, which exhibited similar entry distances.

Across most initial positions, deformability had a smaller effect on target stream entry than cell size or initial position. However, WBC entry distance was strongly dependent on cell deformability for cells initiated at position 1. This observation was consistent with slow migration away from the unstable equilibrium positions at the centres of the short channel walls, particularly for smaller and less deformable cells.

In summary, we note that i) larger and more deformable cells followed HCOs located closer to the channel centre, ii) the overall HCO shape was similar in all cases, iii) larger cells migrate faster and reach the target stream earlier and iv) the effect of deformability on entry length is coupled with cell size; greater deformability increases the entry length for the larger CTC but decreases it for the smaller WBC. These single-cell results established the baseline migration pathways and entry distances used below to determine whether CTC-WBC interactions explain the altered WBC migration observed in mixed suspensions.

### 3.3 Effect of CTC deformability on WBC migration in suspension

Fig. 4 compares WBC migration in three suspension conditions: WBC-only reference case (first row) and WBC suspensions containing a CTC with *La*_CTC_= 10 or 100 (second and third row, respectively). In all cases, *La*_WBC_= 100. The side views show cell migration in the depth direction (*z*-axis, Fig. 4a,4d and 4g), while the cross-sectional views at two downstream locations (*x* = 0.25 and 0.75 cm) show lateral cell migration (Fig. 4b, 4c, 4e, 4f, 4h and 4i). The *z*-component of the corresponding single-cell equilibrium position, *z*_eq_, is included in the side views for comparison.

**Fig. 4.**
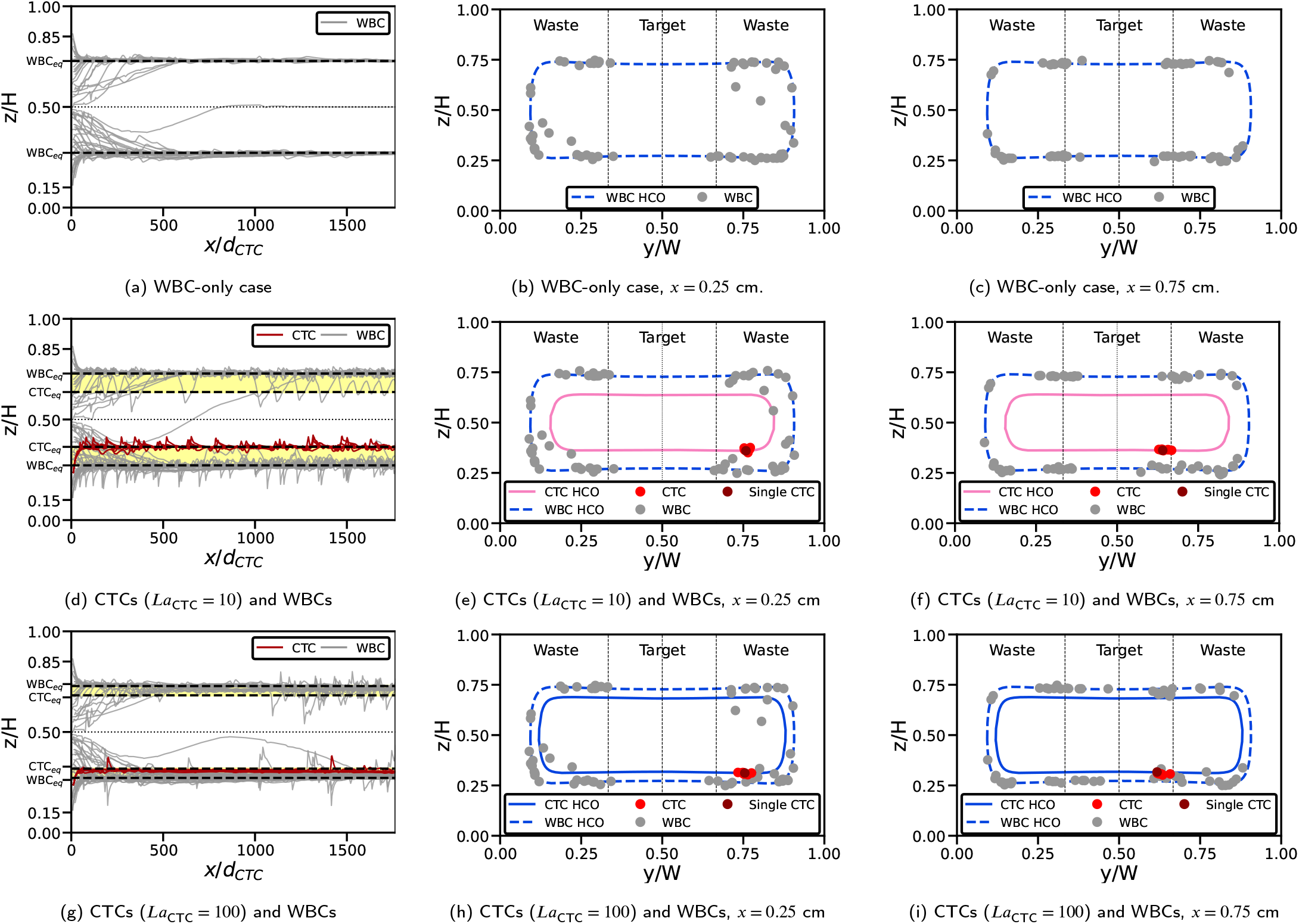
Stacked cell locations from eight separate configurations for three cases: (a–c) WBC-only case; (d–f) CTC with *La*_CTC_ = 10 and WBCs; (g–i) CTC with *La*_CTC_ = 100 and WBCs. In all cases, WBCs have the same deformability (*La*_WBC_ = 100). Panels (a), (d) and (g) show the side-view trajectories, while panels (b), (e) and (h), and (c), (f) and (i), show cross-sectional cell locations at *x* = 0.25 cm and *x* = 0.75 cm, respectively. Corresponding HCOs are shown as lines, and the dark-red marker denotes the position of an isolated CTC.

Fig. 4a shows the WBC-only case, in which all but one WBC migrated to the vicinity of *z*_eq_within a downstream distance of approximately 500*d*_CTC_. Small fluctuations persisted during and after this migration, consistent with interactions among the WBCs.

Fig. 4b and 4c show that the WBCs largely remained close to their single-cell HCO. At *x* = 0.25 cm, the majority of WBCs were located near the HCO, and by *x* = 0.75 cm they had continued to migrate along it.

When a CTC was present in the suspension (Figs. 4d – 4g (second and third row)), some WBCs deviated from their single-cell HCO toward the CTC HCO. When the CTC was more deformable (*La*_CTC_= 10), fewer WBCs deviated from the WBC HCO than when the CTC was less deformable (*La*_CTC_= 100). WBCs also reached the target stream over a shorter downstream distance in the presence of a less deformable CTC than in the presence of a more deformable CTC.

In the suspension containing the CTC with higher deformability (*La*_CTC_= 10, Figs. 4d–4f), both the CTC and WBCs migrated rapidly toward their respective *z*_eq_values and HCOs, and generally remained near these single-cell reference positions. Fluctuations in the trajectories were strongly pronounced but localised, and the cells returned rapidly toward their respective *z*_eq_values.

WBC trajectories were generally more disrupted than the CTC trajectory.

As CTC deformability decreased (increasing *La*_CTC_, Figs. 4g–4i and Figs. A2a–A2c), both the CTC and WBCs migrated to positions slightly away from their respective *z*_eq_values and HCOs, but remained within the region bounded by the CTC and WBCs equilibrium positions. Furthermore, the trajectory fluctuations decrease in intensity and frequency with increasing *La*_CTC_, Figs. (4g, A2a–A2c). The reduced fluctuations were consistent with more persistent CTC-WBC interactions when deformability of the CTC and WBCs was similar ^23^.

WBCs displaced from the isolated-WBC HCO generally migrated toward the target stream over a shorter downstream distance than WBCs in the WBC-only reference case. Additionally, the deviated WBCs were located in the same horizontal channel half as the CTC, and occurred in both the bottom and top regions of the channel. These trajectories suggest that the displaced WBCs underwent long interactions with the CTC in linearly aligned or staggered formations.

In the WBC-only suspension (Fig. 4a), one WBC remained on the midplane. However, when a CTC with *La* ∈ [10, 20, 30, 50] was present, the corresponding WBC migrated into the lateral half opposite the CTC (Figs. (4d,A2a–A2c)). For *La*_CTC_= 100 (Fig. 4g), this WBC instead migrated into the bottom half of the channel with the CTC. This condition-dependent migration suggested that the relative depth positions of the CTC and WBC influenced their interaction dynamics and contributed to the observed dependence of WBC migration on *La*_CTC_.

Overall, these results show that the presence and deformability of a CTC altered WBC migration relative to the single-cell and WBC-only reference trajectories. To determine how these suspension-level differences arose, we next examined how individual CTC-WBC encounters perturbed WBC migration.

### 3.4 Role of CTC-WBC interactions

Fig. 5 shows the migration rates (*y*^′^ = d*y/*d*x*) as a function of the axial centre-to-centre separation distance *δx* from the CTC for *La*_CTC_= 10 (Fig. 5a) and *La*_CTC_= 100 (Fig. 5b). The migration rate was defined as positive (*y*^′^ > 0) when the WBC moved toward the channel centre, irrespective of its quadrant location. The distance was defined as positive (*δx* > 0) when the WBC was downstream of the CTC, and as negative (*δx* < 0) when the WBC was upstream of the CTC. Under the periodic boundary conditions along the *x*-axis, the shortest axial distance between the cells was used, such that |*δx*| < *L/*2. WBC migration rates are segregated according to their quadrant location relative to the CTC as defined in Fig. 1c. WBC migration rates were grouped according to the relative cross-sectional locations of the WBC and CTC, using the quadrants defined earlier. For cells occupying the same lateral half of the channel, scenarios A and B distinguished whether the CTC or WBC, respectively, was closer to the channel center. The WBC migration rates for the WBC-only case are shown in the inset for comparison. For this reference case, *δx* the distance was calculated between each WBC and its nearest axial neighbour. Because no CTC was present, the data were not divided by relative CTC-WBC quadrant. The maximum single-cell migration rates along the HCO and the average migration rates for the WBC-only suspension were included as horizontal lines for reference.

**Fig. 5.**
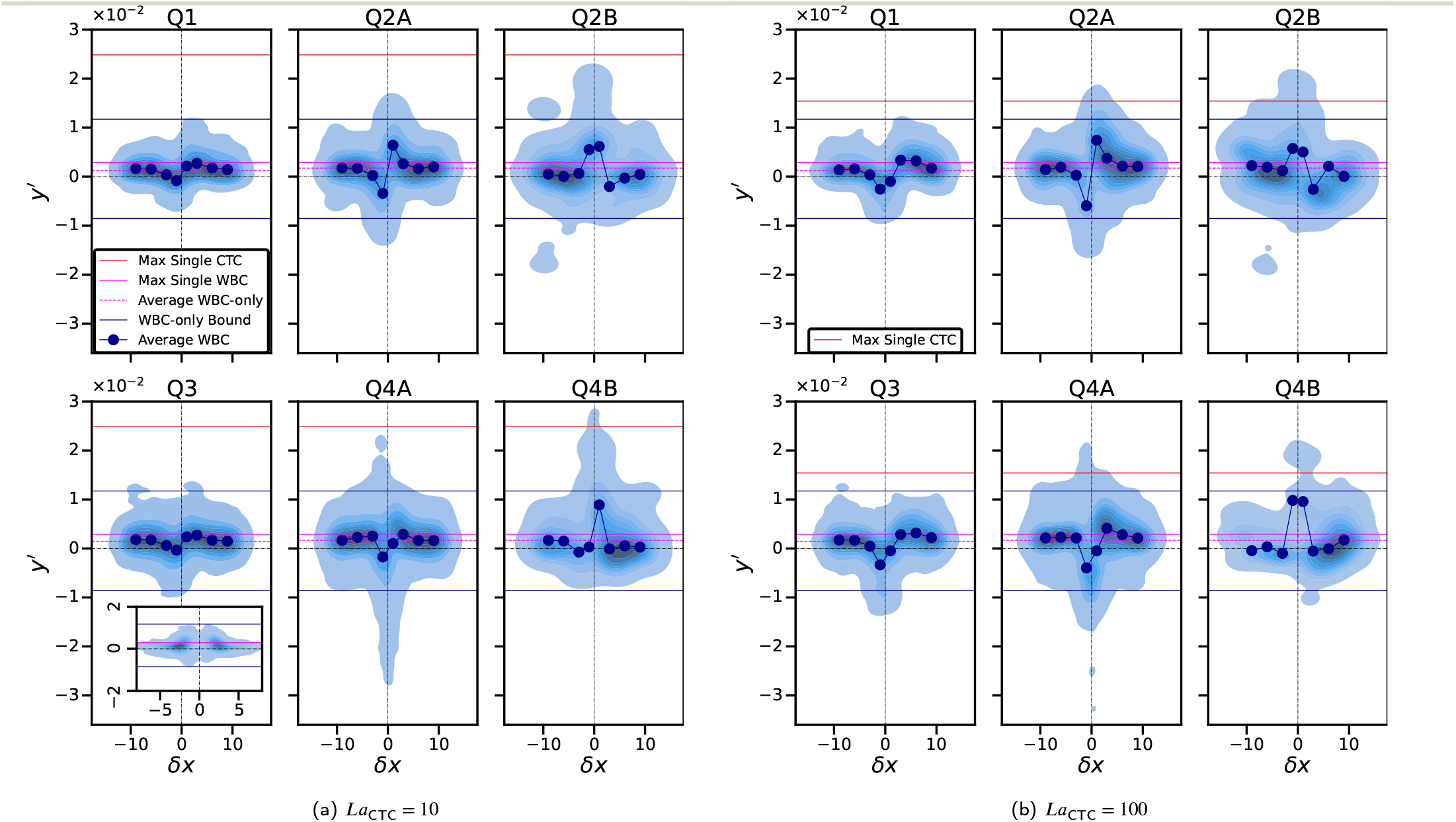
WBC migration rates *y*^′^ = d*y/*d*x* in relation to their axial centre-to-centre distance *δx* to the CTC for (a) *La*_CTC_ = 10 and (b) *La*_CTC_ = 100. The migration rate is defined as positive (*y*^′^ > 0) when the WBC moves toward the channel centre, and the distance is taken as positive (*δx* > 0) when the WBC is leading the CTC. Depending on its location in the channel cross-section with respect to the CTC, the WBC is counted in one of the six distributions (see Fig. 1c). The inset shows the WBC-only case where shortest axial distances between WBCs are used for *δx*. The solid horizontal lines indicate the maximum single CTC and WBC migration rates, as well as the bounds for the WBC-only case. The circles mark the average WBC migration rate for the binned values of *δx*.

In the WBC-only suspension (inset in Fig. 5), the WBC migration rates were largely comparable to those observed for single-cell behaviour. Some variation occurred when a WBC was close to another WBC (|*δx*| < 3*d*_WBC_). This variation included both increases and decreases in the migration rate, in some cases exceeding the magnitude of the maximum single-WBC migration rate. However, the distribution of these variations was concentrated near *δx* ≈ ±2.5*d*_WBC_. This spacing was consistent with the preferred spacing for pair and train formation identified by Kahkeshani *et al*. ^24^, suggesting that nearby WBCs interacted while maintaining axial separation.

In the presence of a CTC (Fig. 5a and Fig. 5b), large distance-dependent variations occurred in WBC migration rate in all quadrants. Two general patterns were observed for both CTC Laplace numbers. For Q1, Q2A, Q3, and Q4A the average value of *y*^′^was close to the single-cell value for large distances (|*δx*| ≈ 10*d*_CTC_). As the cells approached, *y*^′^was reduced, and sometimes became negative, for *δx* just below zero, while *y*^′^ was increased for *δx* just above zero. For Q2B and Q4B, a pronounced increase in *y*^′^ occurred for *δx* ≈ 0.

These patterns depend on the relative positions and orientation of the cells. In Q2A and Q4A, the CTC and WBC occupied the same lateral half of the channel, with the CTC closer to the channel centre. The CTC approached and overtook the WBC as *δx* decreased from positive to negative values. During this interaction, the WBC initially migrated towards the channel center (*y*^′^ > 0) and subsequently migrated toward the wall (*y*^′^ < 0) before resuming center-directed migration after the CTC moved downstream. For Q2B and Q4B, the WBC was closer to the channel centre than the CTC. The WBC moved faster along the channel and overtook the CTC as *δx* increased from negative to positive values. During this interaction, the presence of the CTC on the wall side of the WBC was associated with a pronounced center-directed WBC migration (*y*^′^ > 0). In all of these cases (Q2A, Q2B, Q4A, Q4B), cells approached closely and the magnitude of *y*^′^ sometimes exceeded the maximum migration rate of an isolated WBC by more than ten times. These large perturbations were thus associated with close CTC-WBC proximity. In Q1 and Q3, where the WBC and CTC were on different sides of the channel (*y/W* < 0.5 and *y/W* > 0.5, respectively) and, therefore, remained farther apart. WBC migration rates still differed from the single-WBC values, but the deviations were smaller than those observed when two cells occupied the same half.

Fig. 5 shows that CTC deformability did not change the general trends of WBC migration, but affected the magnitude of the WBC migration rates. Comparing the results for the CTC with the highest deformability investigated (*La*_CTC_= 10) with the CTC with the lowest deformability (*La*_CTC_= 100), the averaged profiles reached more extreme values for the CTC with lower deformability, while individual peak magnitudes were larger for the more deformable CTC. These differences occurred when the WBC was in close proximity to the CTC, suggesting that cell deformability plays a role in the close CTC-WBC interactions.

Overall, Fig. 5 indicates that CTC proximity produced substantial, position-dependent perturbations in WBC migration rate and that these interactions depend on the CTC deformability. However, a large migration rate did not necessarily produce a large next WBC migration because the outcome also depended on the duration of the interaction. Therefore, it is still unclear through which mechanism CTC deformability affects the WBC-CTC interactions.

We therefore examined individual CTC-WBC interactions while the WBCs remained within the waste streams. Using the trajectory analysis described in Section 2.4, the interactions were classified as short or long. Representative trajectories of each type are shown in Fig. 6. A **short interaction** occurred when one cell overtook the other (Fig. 6a). During this interaction, the WBC was displaced either toward the target stream or back towards the wall (Fig. 6b) and temporarily departed from its single-cell HCO (Fig. 6c). The corresponding WBC migration rate was affected for a short time only (Fig. 6d). These short interactions therefore are responsible for the larger instantaneous WBC migration rates observed in Fig. 5, although the direction and net migration varied. A **long interaction** occurred when the CTC-WBC pair maintained a similar velocity and remained in close proximity (Fig. 6e). During this interaction, the WBC moved closer to the CTC (Fig. 6f), departed from the isolated WBC HCO, and migrated within the region between the WBC and CTC HCOs (Fig. 6g). The two cells subsequently migrated at similar rates, approaching that of a single CTC (Fig. 6h). The prolonged interaction produced a greater net WBC migration than a single WBC under the same flow conditions. Long interactions, therefore, are responsible for larger average WBC migration toward the target stream.

**Fig. 6.**
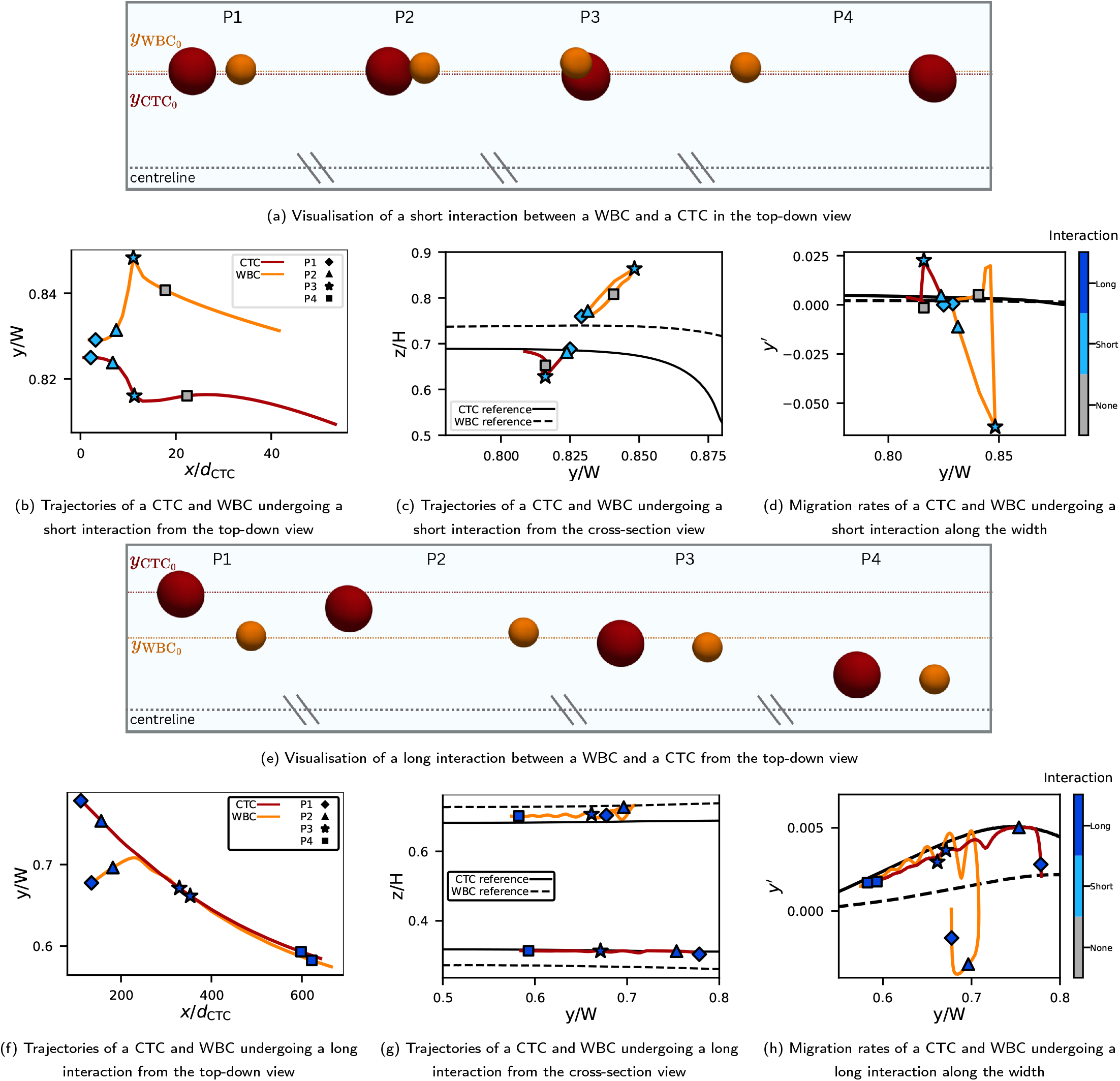
Short and long interactions between a CTC and a WBC. Top-down visualisation of (a) short interaction and (e) long interaction. Trajectories of the migration in the top-down view of (b) short interaction and (f) long interaction. Trajectories of the migration in the cross-section view of (c) short interaction and (g) long interaction. Migration rates along the width of a (d) short interaction and (h) long interaction. Four downstream positions of the short and long interactions were chosen to be visualised in (a) and (e), these positions are represented by markers in (b-d) and (f-h), respectively, and are coloured by the type of interaction at that position.

WBC trajectories that were neither a short nor a long interaction were assigned to the ‘none’ category. This category is representative of single WBC migration, and WBC-WBC interactions, which had little effect on their migration rates (Fig. 5a inset).

Overall, short and long CTC-WBC interactions had distinct migration signatures. Short interactions produced the largest changes in instantaneous WBC migration rate, but the direction and net migration varied. Long interactions produced smaller or less variable changes in WBC migration rate over a longer downstream distance, resulting in greater net WBC migration toward the target stream. We next determined how the occurrence of these interactions depended on the mechanical properties and relative HCOs of the two cell types.

### 3.5 Relative HCO separation affect CTC-WBC interactions

To determine the conditions associated with short and long CTC-WBC interactions, we varied the deformability of both cell types. The CTC and WBCs Laplace numbers were varied independently (*La*_CTC_, *La*_WBC_∈ [10, 30, 100]), resulting in a 3 ×3 parameter space of mechanical conditions. Observations were weighted by the inverse of the number of observations per case so that each case contributed equally to the analysis. Consequently, each of the nine conditions contributed equally to the pooled analysis.

Fig. 7 shows the distribution of the interaction types across the nine mechanical conditions. Long interactions occurred in all quadrants, including configurations in which the cells were longitudinally aligned or staggered across the channel cross-section. Generally, as the deformability ratio *La*_WBC_*/La*_CTC_decreased, the weighted fraction of long-term interactions increased. However, the equal-deformability cases (*La*_WBC_= *La*_CTC_) differ from each other as the weighted fraction of long interactions was greater when both cell types were less deformable (larger *La*). Thus, the deformability ratio alone did not account for the observed distribution of interaction types.

**Fig. 7.**
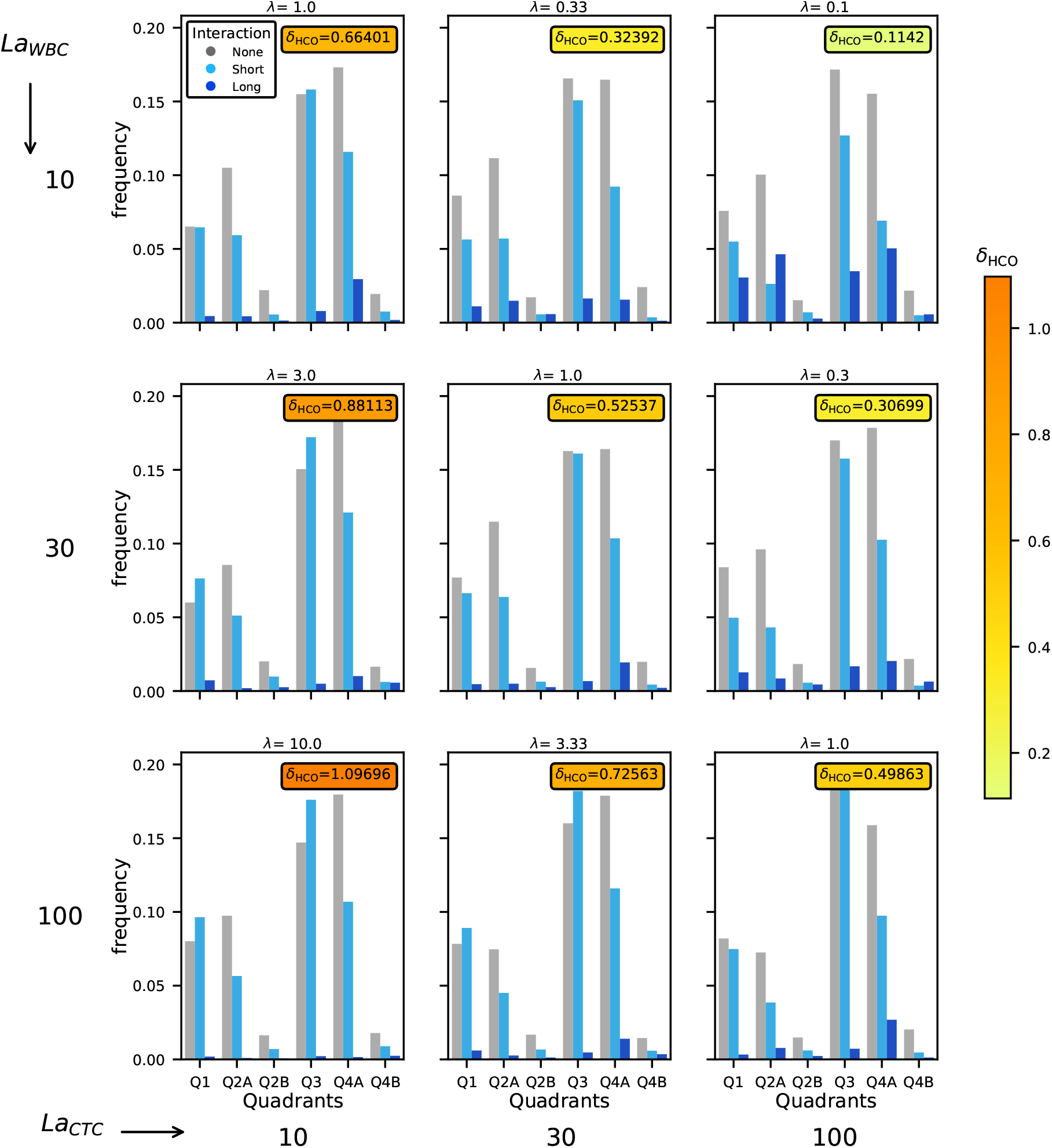
Weighted interaction counts for suspensions with varied deformability ratios *λ*, grouped into four quadrants with two sub-quadrants. CTC deformability decreases from left to right, while WBC deformability decreases from top to bottom. The mean radial distance metric, (D), defined relative to the WBC radius *δ*_HCO_ = D*/*d_WBC_*/*2, is indicated in the upper-right corner of each sub-figure. Values are colour-coded from largest to smallest separation. The deformability ratio, *λ* = *La*_*W BC*_ */La*_*CT C*_, is denoted in the title of each sub-figure.

Previous studies showed that equilibrium positions influence particle pair formation ^25,26^. We therefore examined whether the relative migration trajectories of the two cell types were associated with interaction persistence by comparing the nine mechanical conditions using the mean separation between the singlecell HCOs, *δ*_HCO_. The occurrence of long interactions generally decreases as *δ*_HCO_increased. In particular, the largest distance between HCOs was achieved for the most deformable CTC (*La*_CTC_= 10) and the least deformable WBCs (*La*_WBC_= 100). Under this condition, *δ*_HCO_≈ *r*_WBC_and the lowest weighted fraction of long interactions was observed. Conversely, the smallest *δ*_HCO_occurred for the least deformable CTC (*La*_CTC_= 100) and the most deformable WBC (*La*_WBC_= 10), for which the greatest weighted fraction of long interactions was observed. Cells with closer HCOs experienced similar axial flow velocity. We hypothesise that the smaller relative velocity allowed the cells to remain in close proximity for a longer downstream distance, allowing the two cells to hydrodynamically interact for longer.

Overall, varying cell deformability affected the location of its HCO and thus changed the relative migration trajectory of CTCs and WBCs. Greater separation between the HCOs of the CTC and the WBCs was associated with no or short interaction, while closer

HCOs were associated with a greater fraction of long interactions. In addition to this indirect effect of cell deformability on cell-cell interaction, Owen *et al*. ^23^ demonstrated that cell deformability directly affects the close interaction of pairs with prescribed initial positions. Together, these findings indicate that deformability influences both how cells approach one another and their subsequent interactions. These combined effects alter WBC migration relative to the single-cell trajectory and can promote WBC migration toward the target stream.

### 3.6 Design implications and model limitations

Patient-derived CTCs are heterogeneous in size and mechanical properties ^3,4^, which may alter the relative separation between the cell-specific HCOs of CTCs and WBCs. Our findings suggest that maximising the distance between these HCOs may reduce long CTC-WBC interactions and thus limit WBC migration toward the target stream. However, HCO separation should not be maximised independently because the same operating change may alter HCO engagement, target-stream entry length, or outlet partitioning. Consequently, device optimisation should balance reliable CTC and WBC migration with reduced susceptibility to CTC-WBC interactions.

As a result, single cell simulations can be used to evaluate relative CTC and WBC HCOs across different device geometries and flow conditions, before mixed suspension simulations or experiments are performed. In this context, *δ*_HCO_may be useful as a computational metric for identifying conditions that warrant more detailed mixed-suspension analysis. Potential strategies include adjusting the flow rate to alter HCO shape and location, as demonstrated by Nakagawa *et al*. ^15^, controlling the inlet flow split to alter initial cell positions and HCO engagement, and selecting a channel length that permits CTC migration into the target stream while limiting opportunities for long CTC-WBC interactions.

The present study was limited to the geometry and flow condition of one channel. Although relative HCO separation provides a useful descriptor under these conditions, the relationship between HCO separation and interaction persistence may differ with channel aspect ratio, confinement, flow rate, or inlet configuration. Evaluation across additional operating conditions will be required before *δ*_HCO_can be used as a general device optimisation criterion. Furthermore, *δ*_HCO_is an area-based estimate of the mean separation between isolated-cell HCOs. It does not account for differences in the sections of each HCO or interactions occurring away from the HCO. Thus, *δ*_HCO_is a descriptor of interaction susceptibility rather than a complete predictor of individual CTC-WBC interactions.

We made several modelling assumptions to isolate the effects of the cell mechanical properties and CTC-WBC interactions on inertial migration. Cells were assumed to be spherical elastic capsules, without internal structures, and with a fixed size. This allowed us to study the effect of cell deformability, independently under controlled conditions, but did not reproduce the full mechanical heterogeneity of patient-derived CTCs and WBCs. The investigated Laplace numbers represent selected parametric conditions rather than specific patient-derived cell phenotypes. Consequently, the results establish trends with relative deformability rather than quantitative predictions for a particular biological cell population.

Future studies should investigate the effects of cell size distributions, non-spherical resting shapes, and intracellular structures on migration and CTC-WBC interactions. Cells were initialised at randomly distributed positions when entering the straight channel, and the inlet and outlet bifurcations were omitted to reduce computational cost. Consequently, the simulations did not ac-count for cell migration or CTC-WBC interactions within the inlet transition, nor did they directly resolve the cell fractionation at the outlets. The modelled target stream should therefore be interpreted as a geometric proxy for predicted outlet routing rather than as a direct calculation of WBC carryover or sample purity. Previous experimental studies established the relevance of the device and isolated-cell migration framework, but the specific interaction types identified here were not validated experimentally. The present results therefore provide mechanistic predictions that can guide future measurements of CTC-WBC interactions.

The CTC:WBC ratio was intentionally increased relative to that in clinical samples to provide sufficient CTC-WBC interactions for mechanistic analysis. Although the number of WBCs encountered by a CTC was consistent with that estimated for experimental cell concentration ^14^, the observed interaction frequencies between a CTC and WBC should not be interpreted as expected frequencies in patient blood. Streamwise periodic boundary conditions allowed faster migration CTCs to overtake ‘new’ WBCs as they travel downstream. Thus, a CTC could encounter more WBCs than the eight WBCs explicitly represented within one periodic domain. Although previous validation indicates that cells did not interact with their own periodic images ^14^, periodicity may still influence the sequence and frequency of repeated encounters. The interaction statistics should therefore be interpreted as a comparison among mechanical conditions within the present model rather than as absolute event frequencies for the physical device.

Despite these limitations, the simulations identify a mechanism by which cell mechanical properties alter relative CTC and WBC migration, interaction persistence, and WBC migration from the single-cell trajectory. Collectively with previous work on HCOs, these results indicate that inertial microfluidic separation should be evaluated in terms of single-cell migration rates; equilibrium position; HCO location, size, and engagement; and susceptibility to interactions.

## 4 Conclusions

Inertial microfluidic separation occurs within mechanically heterogeneous cellular suspensions, yet device behaviour is commonly interpreted using isolated-cell migration. Using 3D simulations of an established straight-channel device, we examined how the deformability of CTCs and WBCs affected cell-specific HCOs, interaction types, and WBC migration toward the target stream.

The presence and deformability of a CTC altered WBC migration relative to the WBC-only reference. Short interactions produced large but brief migration-rate changes in either direction, while long interactions maintained CTC-WBC proximity and produced WBC migration toward the target stream. Smaller relative HCO separation was associated with a greater number of long interactions, indicating that deformability affected WBC migration by both changing the relative cell-specific trajectory and modifying the subsequent interaction between the cells. This finding becomes particularly important as early-stage cancer cells are known to exhibit reduced deformability, ^3 4^ which reduces HCO separation, therefore, potentially reducing separation efficiency and compromising the detection of CTCs at an early stage of disease.

Our work shows the complexity of the CTC-WBC separation problem in inertial microfluidics. It highlights that single-cell migration does not fully determine separation in a heterogeneous suspension. Relative HCO geometry provides a mechanistic descriptor of interaction susceptibility that complements equilibrium position and HCO engagement. Increasing the separation between CTC and WBC HCOs may reduce long interactions, whilst maintaining reliable CTC and WBC migration rates, may improve separation efficiency in IMF devices.

## 5 Acknowledgements

The authors acknowledge UK Research and Innovation (UKRI) and the Engineering and Physical Sciences Research Council (EPSRC) for awarding computational resources under the Access to HPC (Autumn 2025) opportunity, utilizing the Cirrus facility.

## 6 Data availability

The data that support the findings of this study are available on request.

## Appendix

**Figure A1.**
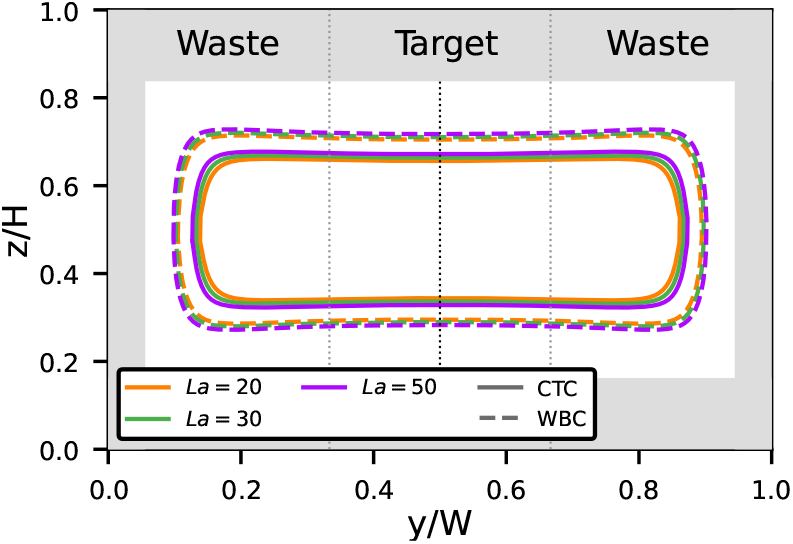
HCOs for the CTC and WBC with softnesses *La* = 20,30,50

**Table A1.** List of physical and numerical simulation parameters.

| Variable | symbol | physical | computational |
| --- | --- | --- | --- |
| Channel width | $W$ | 150 $\mu\text{m}$ | 240 $\Delta x$ |
| Channel height | $H$ | 50 $\mu\text{m}$ | 80 $\Delta x$ |
| Channel length (periodic) | $L$ | 375 $\mu\text{m}$ | 600 $\Delta x$ |
| Hydraulic diameter | $D_h$ | 75 $\mu\text{m}$ | 120 $\Delta x$ |
| CTC diameter | $d_{\text{CTC}}$ | 17 $\mu\text{m}$ | 27.2 $\Delta x$ |
| WBC diameter | $d_{\text{WBC}}$ | 11 $\mu\text{m}$ | 17.6 $\Delta x$ |
| Flow rate | $Q$ | 300 $\mu\text{L}/\text{min}$ | |
| Average flow speed | $\bar{u} = Q/(HW)$ | 2/3 m/s | 0.0417 $\Delta x/\Delta t$ |
| Kinematic viscosity | $\nu$ | $1 \cdot 10^{-6} \text{ m}^2/\text{s}$ | 0.1 $\Delta x^2/\Delta t$ |
| Density | $\rho$ | 1000 $\text{kg}/\text{m}^3$ | 1 |
| Reynolds number | $Re = \bar{u}D_h/\nu$ | | 50 |
| CTC confinement | $\chi_{\text{CTC}} = d_{\text{CTC}}/H$ | | 0.34 |
| WBC confinement | $\chi_{\text{WBC}} = d_{\text{WBC}}/H$ | | 0.22 |
| Grid spacing | $\Delta x$ | 0.625 $\mu\text{m}$ | |
| Time step | $\Delta t$ | 0.039 $\mu\text{s}$ | |
| Relaxation time | $\tau$ | | 0.8 $\Delta t$ |

**Table A2.**
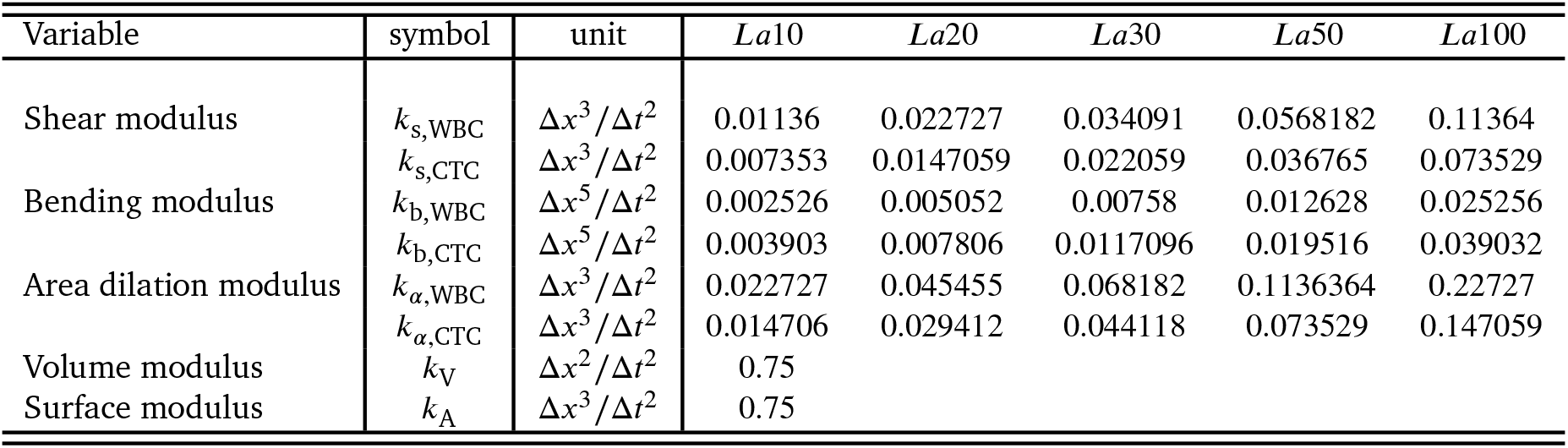
List of numerical parameters varied for CTC and WBC softness.

**Figure A2.**
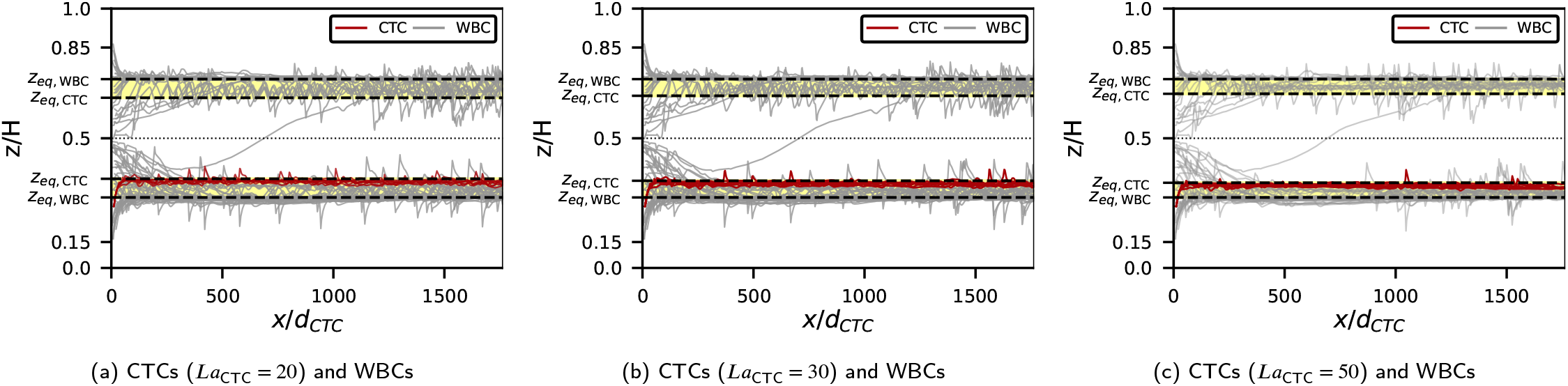
Side view of stacked trajectories from eight separate configurations of six cases: (a) WBC-only case; (b)–(f) CTCs with varying *La*_CTC_ and WBCs. In all cases, WBCs have the same softness (*La*_WBC_ = 100). Horizontal lines indicate corresponding *z*_eq_ values.

**Figure A3.**
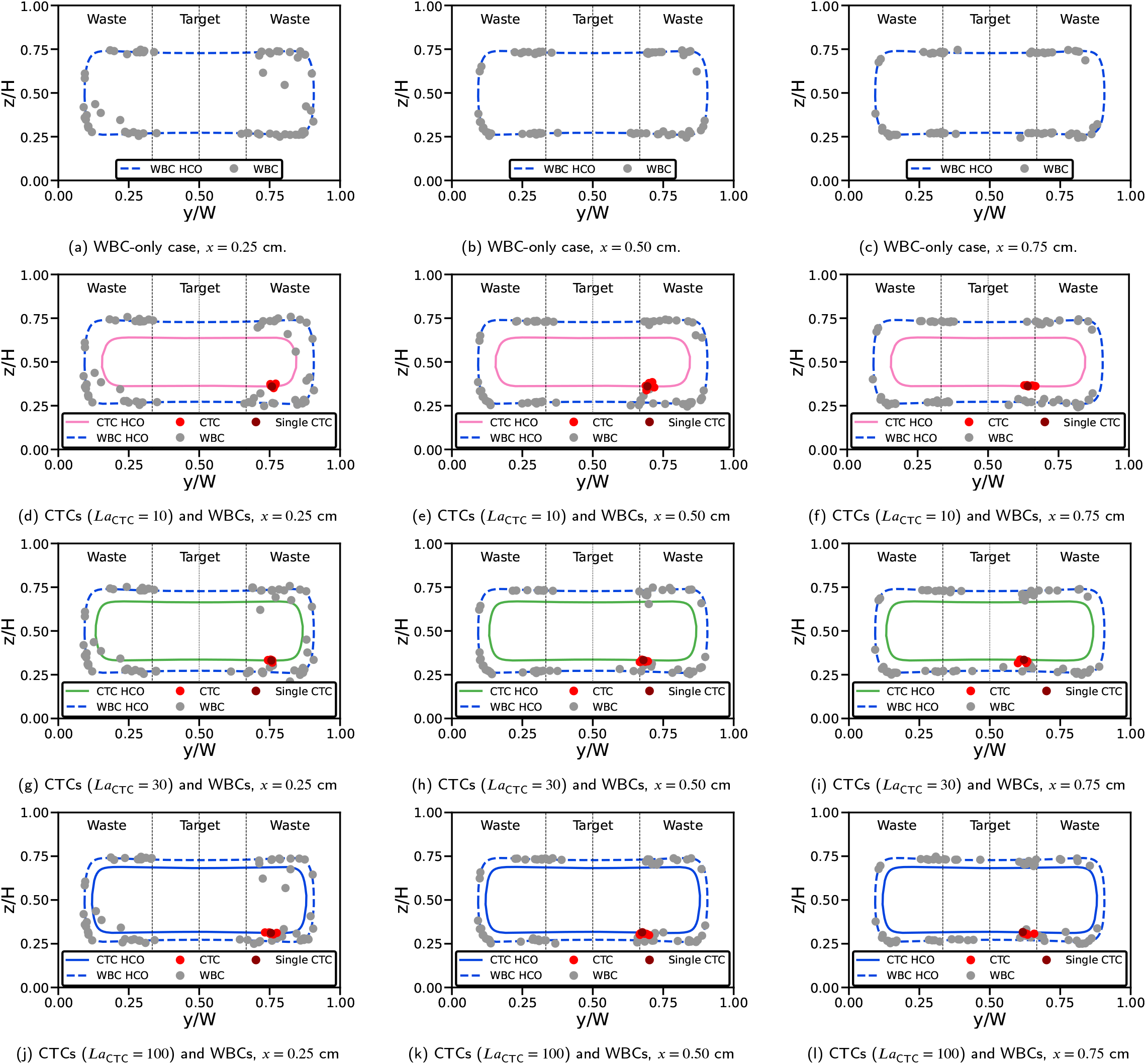
Cross-sectional view of stacked cell locations from eight separate configurations for four cases: (a–c) WBC-only case; (d–f) CTC with *La*_CTC_ = 10 and WBCs; (g–i) CTC with *La*_CTC_ = 30 and WBCs; (j–l) CTC with *La*_CTC_ = 100 and WBCs. In all cases, WBCs have the same softness (*La*_WBC_ = 100). Locations are shown at different downstream locations: *x* = 0.25 cm (first column), *x* = 0.50 cm (middle column), *x* = 0.75 cm (right column). Corresponding HCOs are shown as lines. The dark red dot denotes the behaviour of an isolated CTC.

